# Interplay between periodic stimulation and GABAergic inhibition in striatal network oscillations

**DOI:** 10.1101/073874

**Authors:** Jovana J. Belić, Arvind Kumar, Jeanette Hellgren Kotaleski

## Abstract

The network oscillations are ubiquitous across many brain regions. In the basal ganglia, oscillations are also present at many levels and a wide range of characteristic frequencies have been reported to occur during both health and disease.The striatum is the input nucleus of the basal ganglia that receives massive glutamatergic inputs from the cortex and is highly susceptible to cortical oscillations. However, there is limited knowledge about the exact nature of this routing process and therefore, it is of key importance to understand how time-dependent, periodic external stimuli propagate through the striatal circuitry. Using a large-scale network model of the striatum and corticostriatal projections, here we try to elucidate the importance of specific GABAergic neurons and their interactions in shaping striatal oscillatory activity. Our results show that fast-spiking interneurons, despite their uncorrelated firing, might have a crucial role in the emergence of high-frequency oscillations in the medium spiny neuron population, even if their activity is kept low. Rather, what matters is the firing time relative to just a few other neurons within an oscillation cycle. Finally, we show how the state of ongoing activity, the strengths of different types of inhibitions, the density of outgoing projections, and the overall activity of striatal cells influence network activity. These results suggest that the propagation of oscillatory inputs into the medium spiny neuron population is efficient, if indirect, through fast-spiking interneurons. Therefore, pharmaceuticals that target fast-spiking interneurons may provide a novel treatment for regaining the spectral characteristics of striatal activity that correspond to the healthy state.

**Author Summary:** The striatum is the largest and primary gateway component of the BG, receiving glutamatergic inputs from all cortical areas and is highly susceptible to cortical oscillations. However, there is limited knowledge about the exact nature of this routing process and therefore, it is of key importance to understand how time-dependent, external stimuli propagate through the striatal circuitry. The vast majority of striatal neurons, at least 95% of them, are medium spiny neurons (MSNs) that are also the only source of output from the nucleus. Two of the most examined sources of GABAergic inhibition into MSNs are the feedback inhibition (FB) from the axon collaterals of the MSNs themselves, and the feedforward inhibition (FF) via the small population of fast-spiking interneurons (FSIs) comprising roughly 1-2% of striatal neurons. Using a large-scale network model of the striatum we systematically investigate the propagation of asynchronous periodic cortical inputs, throughout the physiologically relevant range, onto the striatal neurons and their influence on striatal network dynamics. Our results show that FSIs, despite their firing being uncorrelated, may play a crucial role in the efficient propagation of external oscillations onto MSNs. Finally, we show how the state of ongoing activity, the strengths of different types of inhibitions, the density of outgoing projections, and the overall activity of striatal cells influence network activity.

## 1. Introduction

The basal ganglia (BG) comprise the largest subcortical system in the brain thought to be involved in motor control and cognitive processes [1–5]. Dysfunction of neural pathways between cortex and the BG circuitry is linked to many motor disorders, such as Parkinson’s disease (PD), L-DOPA-induced dyskinesia, Tourette’s syndrome, and Huntington’s disease [6–21]. The striatum is the largest and primary gateway component of the BG, receiving glutamatergic inputs from all cortical areas [22]. The vast majority of striatal neurons, at least 95% of them, are medium spiny neurons (MSNs) that are also the only source of output from the nucleus [23–24]. Two of the most examined sources of GABAergic inhibition into MSNs are the feedback inhibition (FB) from the axon collaterals of the MSNs themselves, and the feedforward inhibition (FF) via the small population (1-2% of striatal neurons) of fast-spiking interneurons (FSIs) [25–28]. Feedforward inhibition is powerful, with spiking in a single interneuron capable of significantly delaying spike generation in a large number of postsynaptic spiny neurons, whereas feedback inhibition is made up of many weak inputs [29–35]. High firing rate and uncorrelated spiking of FSIs further amplifies the effect of feedforward inhibition on the MSNs [36–37].

Recently, oscillations (20–80 Hz) have been observed at the level of individual MSNs firing and local field potential in the striatum of both awake and anaesthetized animals [21, 38–40]. The mechanisms underlying the emergence of these oscillations remains unclear. Purely inhibitory network such as the striatum can exhibit network oscillations but this requires very strong input excitation and strong recurrent inhibition [41]. However, given the weak recurrent inhibition [42] in the striatum it seems unlikely that the experimentally observed oscillations are locally generated by the inhibitory network of the striatum. A second possibility is that striatal neurons have sub-threshold resonance properties and when the network is driven strongly enough the oscillations become apparent. However, MSNs do not show any resonance behavior [43] and mechanisms by which resonance behavior of other interneurons can induce oscillations in the MSNs are not understood. A third possibility is that the experimentally observed oscillations in the striatum are in fact cortical oscillations transmitted by the cortico-striatal projections. While the transmission of firing rates and correlations via the cortico-striatal projections has been investigated previously [44–45] it is not clear to what extent cortico-striatal projections can transfer oscillations over a wide range of frequencies and what role do the FB and FF inhibitions play in this process.

Here, we use a biologically realistic, large-scale network model of the striatum to isolate the mechanisms that underlie the transmission of cortical oscillations to not only the MSNs that receive direct cortical input but also to the other unstimulated MSNs. We systematically studied the effect of FB and FF inhibitions, the state of ongoing activity, the density of outgoing projections of FSIs and critically, the overall activity of striatal cells on the oscillatory activity of the striatum. We show that FSIs, despite their uncorrelated spiking, play a crucial role in the efficient propagation of cortical oscillations to the MSNs, especially the ones that did not receive cortical inputs. In the absence of FSI input, spread of oscillations to the unstimulated MSNs requires that either baseline firing rate of the MSNs is unphysiologically large or a very large fraction of MSNs receive the periodic input from the cortex. These findings reveal a new function of FSIs in modulating the transfer of information from the cortex to striatum and by modulating the activity and properties of the FSIs striatal oscillations can be controlled very efficiently.

## 2. Materials and Methods

For the purpose of clarity we firstly describe the model and explain the reasoning behind the choices made in its design.

### 2.1 Neuron models

The neurons in the network were modeled as leaky integrate-and-fire neurons. Some neuron models, such as resonate-and-fire [46] and properly parameterized Hodgkin-Huxley [47], can have different interactions with oscillatory depolarizing currents. The leaky integrate-and-fire neuron model lacks oscillatory features and, therefore, all frequency responses are a direct outcome of network interactions [48]. We considered two types of striatal neurons: medium spiny neurons and fast-spiking neurons. Subthreshold dynamics of the membrane potential *V_i_^MSN^* (*t*) of a MSN *i* was described by the following equation:

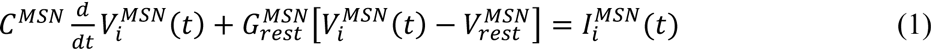

where 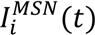 was the total synaptic input current to the neuron, and *C^MSN^* and 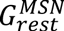 denoted the passive electrical cell properties, the capacitance and conductance of its membrane at rest 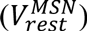, respectively. When the membrane potential reached a fixed spiking threshold *V^MSN^_th_*, a spike was emitted, and the membrane potential was reset to the resting value.

The subthreshold dynamics of the membrane potential 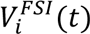 of a FSI *i* was described similarly by the following equation:

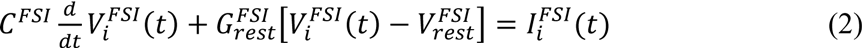

The initial membrane potentials of both MSNs and FSIs were chosen from a uniform distribution from −86.3 to −55 mV and from −82 mV to −65 mV, respectively, in order to avoid any unwanted synchrony, caused by the initial conditions in the simulation runs.

### 2.2 Network

It has been estimated that there are around 2800 MSNs located within the volume of the dendrites of one spiny cell. MSNs are the dominant neuron type in the striatum (up to 95% in rodents [49]), and the radius of their axonal and dendritic arborisations are both around 200 μm. There are at least two inhibitory circuits in the striatum that are activated by cortical inputs and that control firing in MSNs. The first is FF inhibition via the small population of FSIs, and the second is FB inhibition from the axon collaterals of the MSNs themselves. Therefore, we simulated a network of two types of GABAergic neurons, 2800 MSNs and 56 FSIs. A scheme of the striatal network model is shown in Fig. 1A. The connection probability between the MSNs was equal to 0.18 [50]. Each MSN received inhibitory inputs from on average 11 FSIs, resulting in a 20% connectivity from FSIs to MSNs [31].

**Figure 1:**
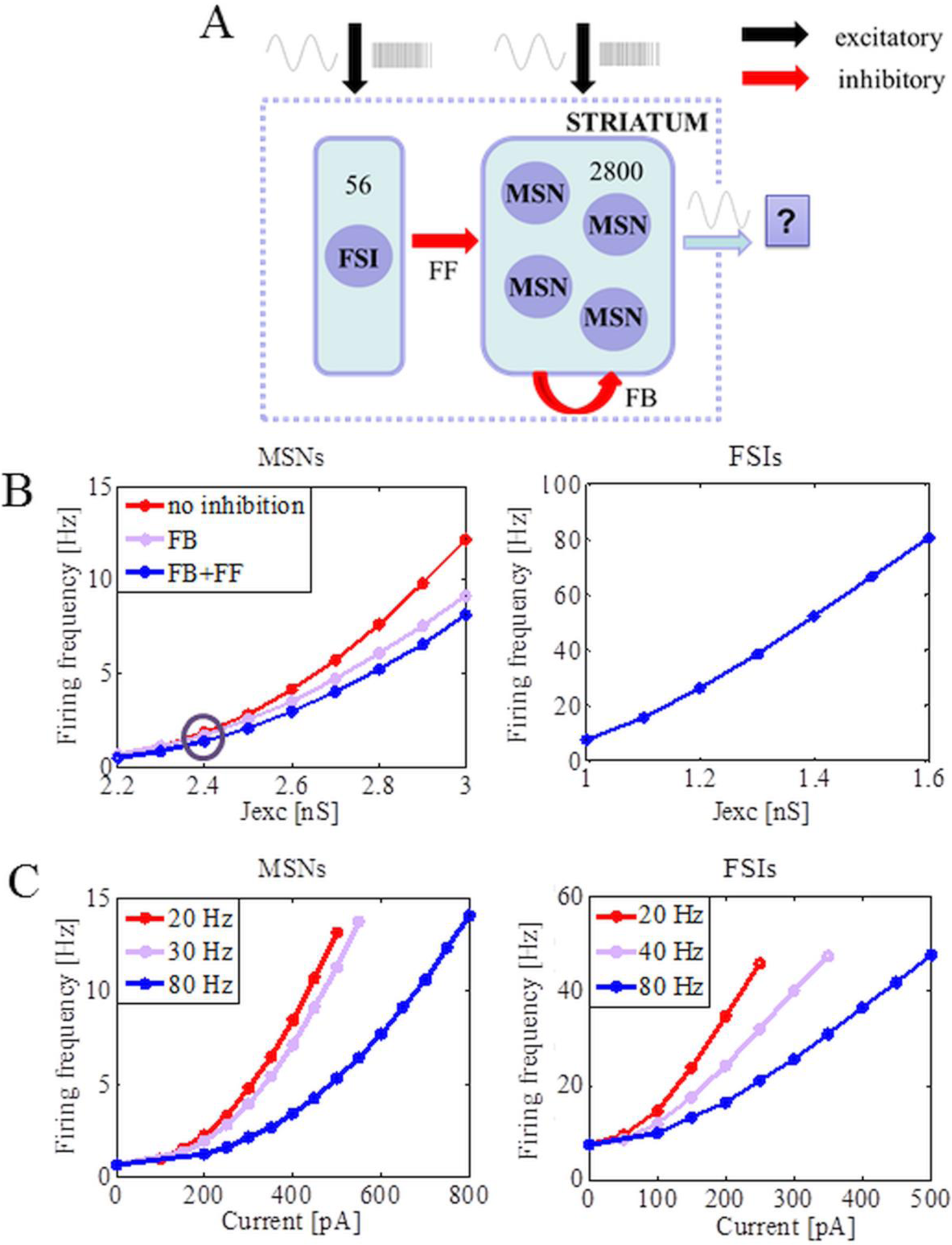
Control of MSNs and FSIs firing rates by external inputs. (A) Schematic of network model with MSNs receiving inhibitory input from FSIs and other MSNs. (B) Excitatory input to both MSNs and FSIs is provided by simulated Poisson trains and calculated by averaging across the neurons within a class (mean ± std). Left panel shows transfer function of MSNs as part of the network with cortical input successfully increased in three cases: without inhibition, when only keeping FB inhibition, when keeping both, FB and FF inhibitions (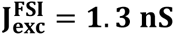 in this case, leading FSIs to fire on average 38.5 ± 6.4 Hz). The circle marks the most frequently used 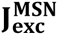. Right panel shows the average firing rate and standard deviation of the firing of FSIs with cortical input successfully increased. (C) Observations of average firing of the connected MSNs and FSIs to oscillatory current injection with different amplitudes and driving frequencies. Additionally, both MSNs and FSIs received simulated background activity that constrains MSNs to fire < 1 Hz (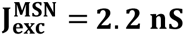) and FSIs <10Hz (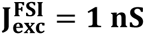).

### 2.3 Synapses

Excitatory synaptic input was modeled by transient conductance changes using the alpha-function such that:

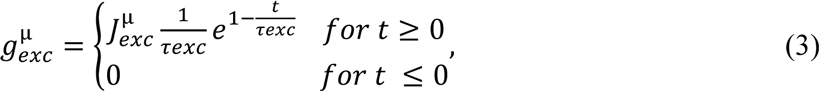

where μ ∊ {MSN, FSI}, *τ_exc_* and 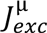 denoted the rise times for the excitatory synaptic inputs and the peak amplitude of the conductance transient (‘strength’ of the synapses), respectively (Fig. 1B). Inhibitory synaptic input was modeled similarly:

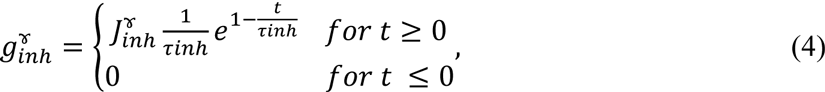

where *γ* ∊ {FF, FB}. The total excitatory conductance 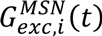 in a MSN *i* was equal to:

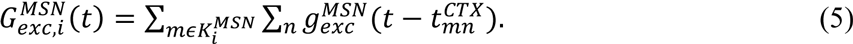

Here, 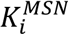 denoted the set of excitatory synapses projecting onto MSN *i*. The inner sum run over the sequence of spikes (*n*’s), and the set 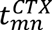 represented the spike times of the excitatory neuron *m*. The total inhibitory conductance 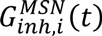 in a MSN *i* was given by:

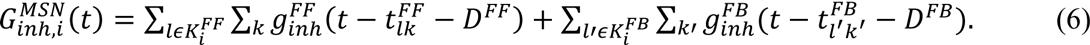

MSNs mainly project onto the dendrites of other MSNs and FSIs tend to project onto the soma, therefore we used a shorter delay for FF inhibition (*D^FF^* = 1 *ms*) compared to the delay used for FB inhibition (*D^FB^ = 2 ms*). Similarly, 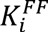 and 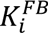 were the sets of presynaptic FSIs and MSNs projecting to MSN *i*. The excitatory conductance 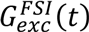 in a FSI *i* was given by:

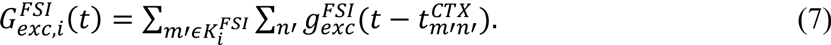

The total synaptic current onto a MSN *i* was:

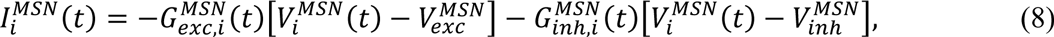

and the total synaptic current onto a FSI *i* was:

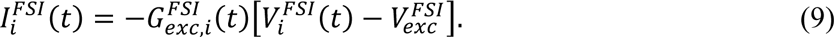

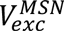 and 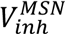 were the reversal potentials of the excitatory and inhibitory synaptic currents of MSNs, respectively, and similarly 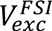 denoted the reversal potentials of the excitatory currents of FSIs.

Based on experimental results, the corresponding strengths of the inhibitory synapses for FF and FB inhibitions in our model, 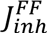 and 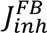, were set to −3 nS and −0.5 nS, respectively, although we tuned the strength of the connections over a range around the experimental values to investigate their influence [31,51]. The parameter values for both MSN and FSIs in our network model are summarized in Table 1.

**Table 1.** Parameter values of the model neurons used in this study.

| Quantity | MSN | FSI |
| --- | --- | --- |
| C [pF] | 120 [34] | 100 [34] |
| $V_{rest}$ [mV] | -86.3 [51] | -82 [34, 52] |
| $V_t$ [mV] | -43.75 [51] | -55 [53] |
| $V_{exc}^\mu$ [mV] | 0 | 0 |
| $V_{inh}^\mu$ [mV] | -65 [49, 54] | -75 [53] |
| $\tau_{exc}$ [ms] | 0.3 | 0.3 |
| $\tau_{inh}$ [ms] | 2 | 2 |
| $G_{rest}$ [nS] | 15.175 [51,55] | 10 [54] |

### 2.4 External input

The selected population of FSIs and MSNs received sinusoidal current, corresponding to an external stimulation from other brain areas, given for neuron *i* as:

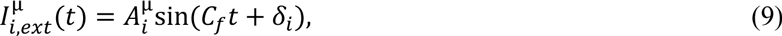

where μ ∊ {MSN, FSI}. The amplitude of oscillations 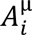 depended on the type of neuron and the driving frequency *C_f_*. It was always set in the 0.9 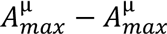 range so that the randomness could be imposed (Fig. 1C). The sinusoidal input was present from the start of the simulation, and δ was a random starting phase in the 0° – 180° range. The rest of population received external, excitatory and uncorrelated Poisson synaptic inputs set to 600 *Hz* to achieve the different realistic firing rates of MSNs and FSIs (Fig. 1B). The background activity constrained MSNs to fire < 1 Hz and FSIs <10Hz.

### 2.5 Implementation

All network simulations were written in the Python interface to NEST. The dynamical equations were integrated at a fixed temporal resolution of 0.01 *ms* using the fourth order Runge-Kutta method. This small time step was necessary because of the fast dynamics of the FSIs. Stimulation ran for 1000 *ms* and each explored scenario was repeated and averaged 10 times, yielding 10 trials. We assumed that plasticity had no significant impact on the network dynamics due to the short duration of stimulation.

### 2.6 Model limitations

Neuronal and network oscillatory dynamics are ubiquitous in real systems, although some recent studies showed that MSNs’ frequency preference did not have a fixed frequency and that MSNs also show no input impedance resonance [43]. In this study, we wanted to better understand the sole striatal network response to oscillatory input and therefore we reduced our model in order to tackle this complex question. The 2800 MSNs located within the volume of the dendrites of one spiny cell constitute only a small volume of striatum and, hence, it is reasonable to assume a distance-independent random connectivity in the network. It is worth mentioning that our results remained qualitatively unchanged when we assumed distance-dependent connectivity (data not shown). In the striatum, FSIs are interconnected by gap junctions, but it is showed that the synchronization effects due to gap junctions are low [56]. Modulatory effects of tonically active and dopaminergic neurons were taken into account by changing the effective strength of the FB and FF inhibitions based on experimental data. Persistent low-threshold spiking (PLTS) neurons are also known to inhibit MSNs, but their output is relatively weak and sparse, therefore inclusion of these neurons would not affect our conclusions [34].

### 2.7 Data analysis

The firing rate of individual neurons was estimated as the average spike count per second. The mean network firing rate was then obtained by averaging the firing rates of all neurons in the network. Network activity in the population that consisted of MSNs was further evaluated by counting the overall spiking activity in time bins of 5 ms, which was practically equal to the number of active cells per time bin.

To estimate the strength of oscillations we used the fact that oscillations introduce peaks in the power spectral density of the population activity. Therefore, we estimated the spectrum *S_pop_* (*f*) of the population activity, directly from the spike trains, using the Fast-Fourier-Transform with the sampling frequency set to 200 Hz. We defined the oscillations index as the relative power in the frequency band of interest as (±5 Hz around the driving frequency *C_f_*):

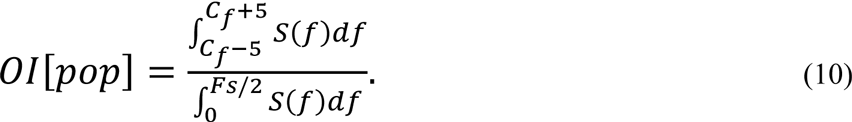

When the network was oscillating strongly, most of the power was contained in the *C_f_* ± 5 band and, as a result, *OI* values were higher.

To measure the spike train similarity, we used Pearson’s correlation coefficient:

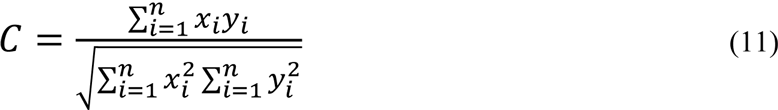

where *x_i_* and *y_i_* denoted the two zero-mean vectors of two spike trains. For identical vectors *x_i_* and *y_i_*, the correlation C was equal to unity. Since we were interested in millisecond synchronization, the bin size was set to 1 ms.

The values in different groups were compared using the Mann-Whitney U-test and a p < 0.01 was considered statistically significant.

## 3. Results

The striatum is a recurrent inhibitory network driven by excitatory inputs from the cortex, where the FF inhibition from FSIs to MSNs and FB inhibition from MSNs to other MSNs are clearly segregated. Individual striatal neurons do not act independently to affect rhythmic population activity, and here we study how they interact with each other and time-dependent external stimuli to induce oscillatory activity in the MSN population.

### 3.1 Propagation of cortical oscillations in a striatal network

When a small fraction of MSNs were stimulated with sinusoidal inputs at 80 Hz (Amax=250 pA) in the addition to the background input, while the rest of MSNs received only excitatory Poisson inputs and fired at 1.74 ± 1.34 Hz, the averaged activity of the MSNs followed the sinusoidal input (Fig. 2, middle, red line). A detailed analysis however reveals that the input driven oscillatory activity was limited only to the stimulated MSNs (Fig. 2, left, red line) and unstimulated neurons did not show any oscillatory activity (Fig. 2, right, red line). That is, partial stimulation of MSNs population is not sufficient to distribute the input to the unstimulated MSNs. By contrast when FSIs were stimulated with the oscillatory inputs (80Hz) oscillations were observed not only in the FSIs but also in all the MSNs (0.04 vs 0.52, p<0.01, Fig. 2, middle; Fig. 2, left). Further we wanted to study sole influence of FF inhibition on the entrainment of the MSNs activity. The analysis revealed that the emergence of a spectral peak in the MSN activity was in fact directly dependent on the average firing of FSIs. For instance, at an 20 Hz input frequency, the FSIs average firing frequency of 14.5 ± 3.3 Hz could not influence activity in the MSN population and induce the peak in the power spectrum, while the average firing at 34.78 ± 4.74 Hz could easily induce a very strong peak (Fig. 3A). Similar results were obtained for 40 Hz and 80 Hz input frequencies in the case of low (red curve) and high (blue curve) firing rate of FSIs (Fig. 3A). Previous experimental work [21, 40, 57] has demonstrated the importance of 80 Hz corticostriatal oscillations and, recently, it has been found that the FSIs fire in relation to very high cortical frequencies [43]. In the following we describe conditions under which an 80 Hz oscillatory input can be transferred to MSN activity. Because FSIs are not connected and receive oscillatory inputs at different phases, they exhibit unstructured activity irrespective of the amplitude of oscillatory input (Fig. 3B, left and middle). Raster plot and firing rate histogram of MSNs, when FSIs fired at high frequencies, also did not show any clear temporal pattern (Fig. 3B, right). Two different values of maximal input oscillatory amplitude corresponded to the average firing of FSIs at 13.21 ± 3.69 and 25.6 ± 4.1, respectively, while the average firing of MSNs did not change drastically 1.62 ± 1.31 Hz versus 1.53 ± 1.26 Hz. Next we wanted to see how frequency and amplitude vary over time (Fig 4B). For A_max_=350 pA the clear temporal oscillatory pattern was present, but the power was not constant over time. The oscillation amplitude and stability of the oscillatory pattern increased with increased A_max_. When the maximal input oscillatory amplitude was set to 250 pA and induced the average firing of FSIs at 21.1 ± 4.25 Hz, the peak at 80 Hz was clearly visible in both cases, for ongoing activity (1.56 Hz; Fig. 4B, left panel) and evoked activity (8.59 Hz, Fig 4B, right panel) in MSNs, although it had a slightly higher value in the second case (0.0357 versus 0.0392, p=0.019).

**Figure 2:**
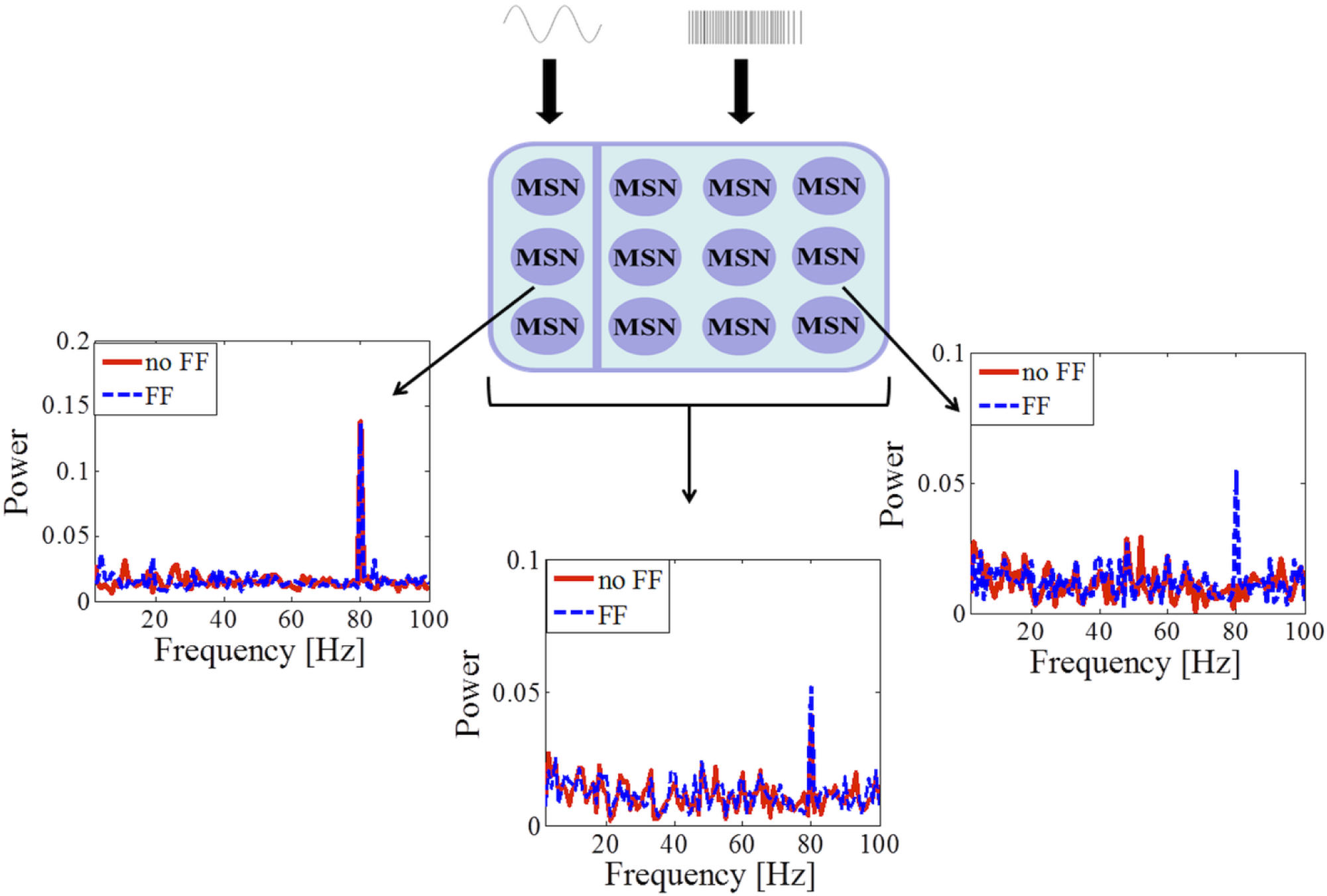
Interplay between oscillatory inputs into MSNs and FSIs. The spectral analysis of the MSN populations where the portion of MSNs receive excitatory inputs from sinusoidal perturbations at 80 Hz and background activity, while the rest of MSNs receive only Poisson input. The analysis was done for the oscillatory, non-oscillatory and whole MSN population in two cases: without FSIs and with oscillatory driven FSIs.

**Figure 3:**
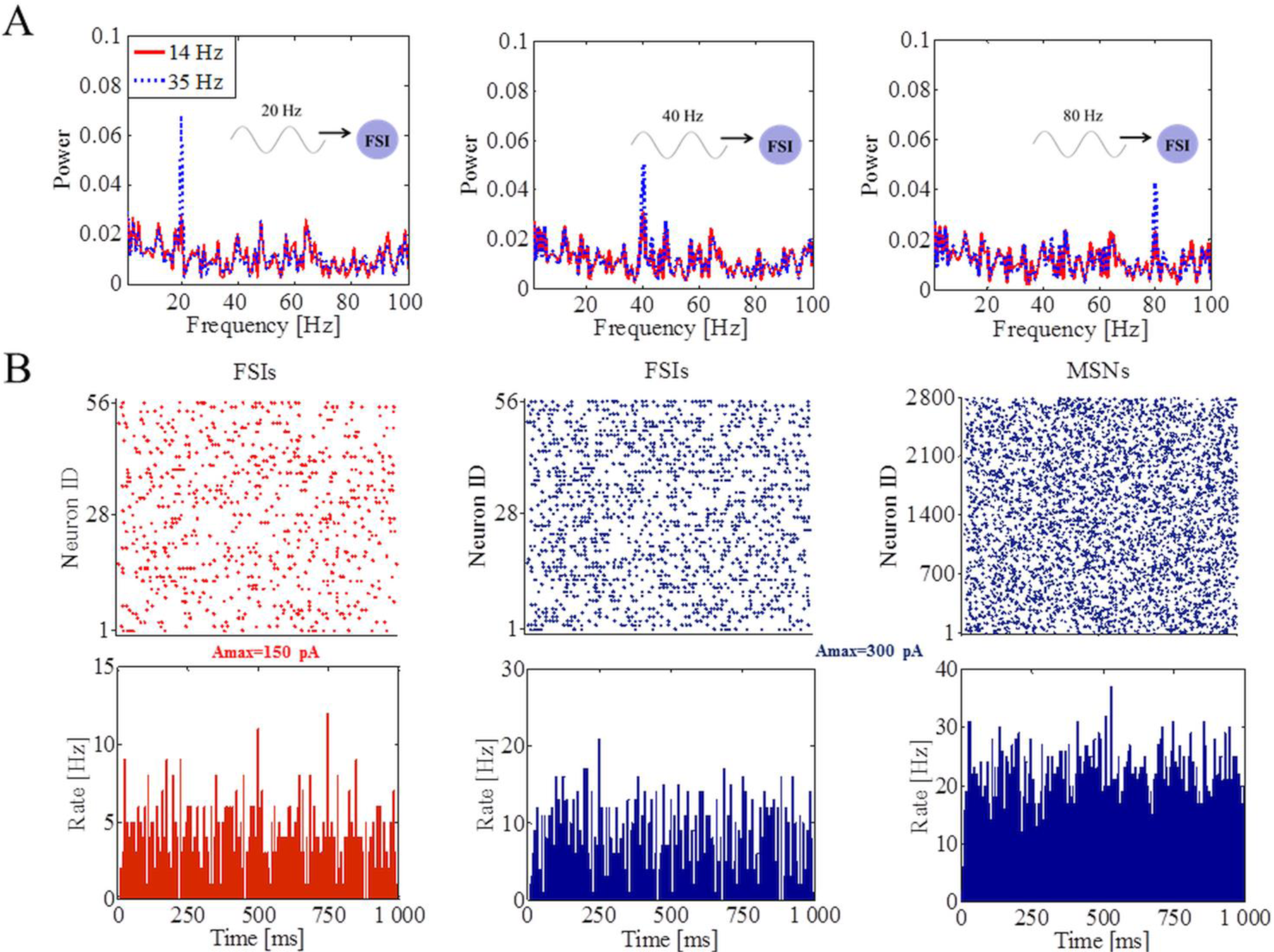
Propagation of asynchronous cortical oscillations in a striatal network model through FSIs. (A) Power spectra of the MSNs population in response to fast cortical oscillatory inputs onto FSIs. Stimulation frequency and amplitude were systematically varied while the network response was analyzed. (B) Raster plots of 56 FSIs for the low (A_max_=150 pA) and high (A_max_=300 pA) maximal oscillatory amplitude onto FSIs (left and middle panel). Raster plot and firing rate histogram of 2800 MSNs for high maximal oscillatory amplitude onto FSIs (right panel).

**Figure 4:**
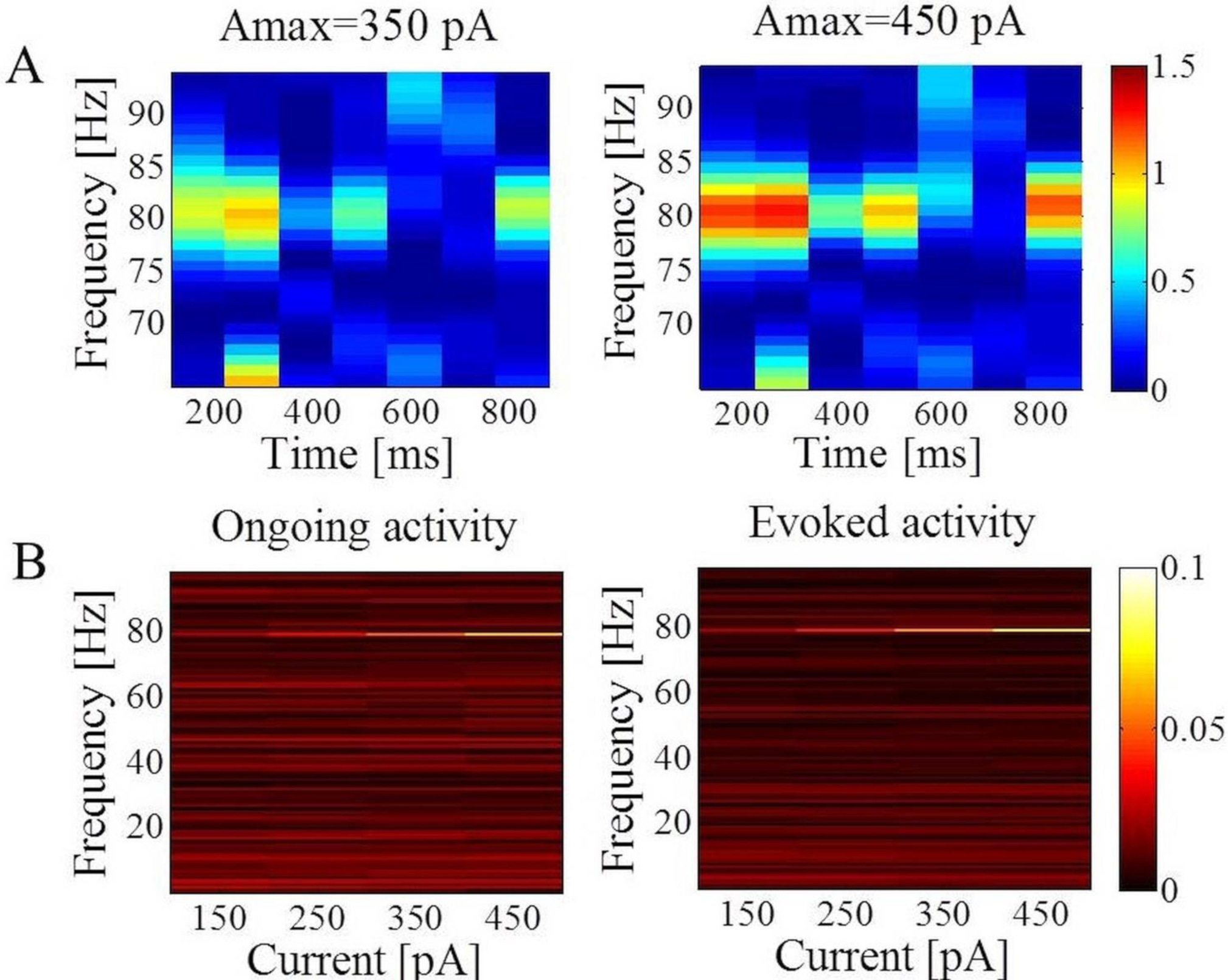
Spectrograms of the striatal MSNs population activity and oscillatory activity in MSN network in response to different average firing values of the MSNs. (A) In the presence of the weak oscillatory input onto FSIs, the network exhibited asynchronous activity. A mild increase in the oscillatory input induced inconstant oscillatory activity in the striatum. The stability and strength of oscillations increased with higher values of A_max_. Colors in the spectrograms indicate the power of oscillation. (B) Power spectra of the MSNs population for different values of the maximal oscillatory amplitude onto FSIs when MSNs underwent ongoing (1.56 Hz; left panel) or evoked (8.59 Hz; right panel).

### 3.2 Firing of FSIs controls propagations of cortical oscillations in the striatal network

The brain needs a mechanism to control the reliable propagation of activity over multiple distant areas. Our computational model showed that FSIs were capable of transferring cortical oscillatory inputs to the MSN population. Excitatory oscillatory inputs to the FSI population (Fig. 5A, left panel) drove the FSIs to fire in a sequential manner. In the cortex and hippocampus, it has been believed that network oscillations require regular spike generation in the population of neurons that form the circuitry [58]. In the striatum, our results showed that just a few FSIs that fire relative to the oscillation cycle were sufficient to transfer periodic fluctuations to the MSN population (Fig. 5A, right panel). Each individual oscillatory cycle spanned over 12.5 ms. The average firing of FSIs was also very low per oscillation cycle (< 0.2 Hz per cycle when the peak in the MSNs started to be prominent; red bars) but increased with a rise in the maximal amplitude of the oscillatory inputs. Each FSI had its own preferred firing time relative to other FSIs. The proportion of neurons (blue bars) that fired in relation to the individual peaks of each FSI separately (defined in a range of over ±2 ms of the maximal value during each cycle) also increased with an increase in the maximal input oscillatory amplitude, but their ratio to the number of neurons that were entrained to the particular cycle declined. To further characterize the activity of the MSN population, we calculated the oscillatory index (OI) and systematically varied the maximal amplitude of the oscillatory inputs (Fig. 5B). The oscillatory index of the MSN population, defined as an integral over power in the range of ±5 Hz around the driving frequency, showed a steady increase correlated with the higher values of the maximal oscillatory input current (for A_max_=150 pA, the peak was not visible, while for A_max_=250 pA, it was quite prominent). Next, we calculated the average pairwise correlation of spiking activity for FSIs. It has been experimentally shown that the firing of striatal FSIs is uncorrelated [36]. Correlation values were very low but increased with an increase in the maximal amplitude of the oscillatory inputs (Fig 5C). The peak was visible in the MSN population even for an average value less than 0.01 for the pairwise correlation. From the perspective of single MSN that receive Poisson excitatory (mimicking external inputs) and inhibitory (mimicking inputs from other neurons) drives, voltage is considerably reduced when oscillatory driven FSIs are added (Fig. 5D). In order to successfully transfer oscillations into MSNs, it is necessary that the number of FSIs that fire during each cycle cross a threshold necessary to modulate activity of sufficient number of MSN within the whole MSN population.

**Figure 5:**
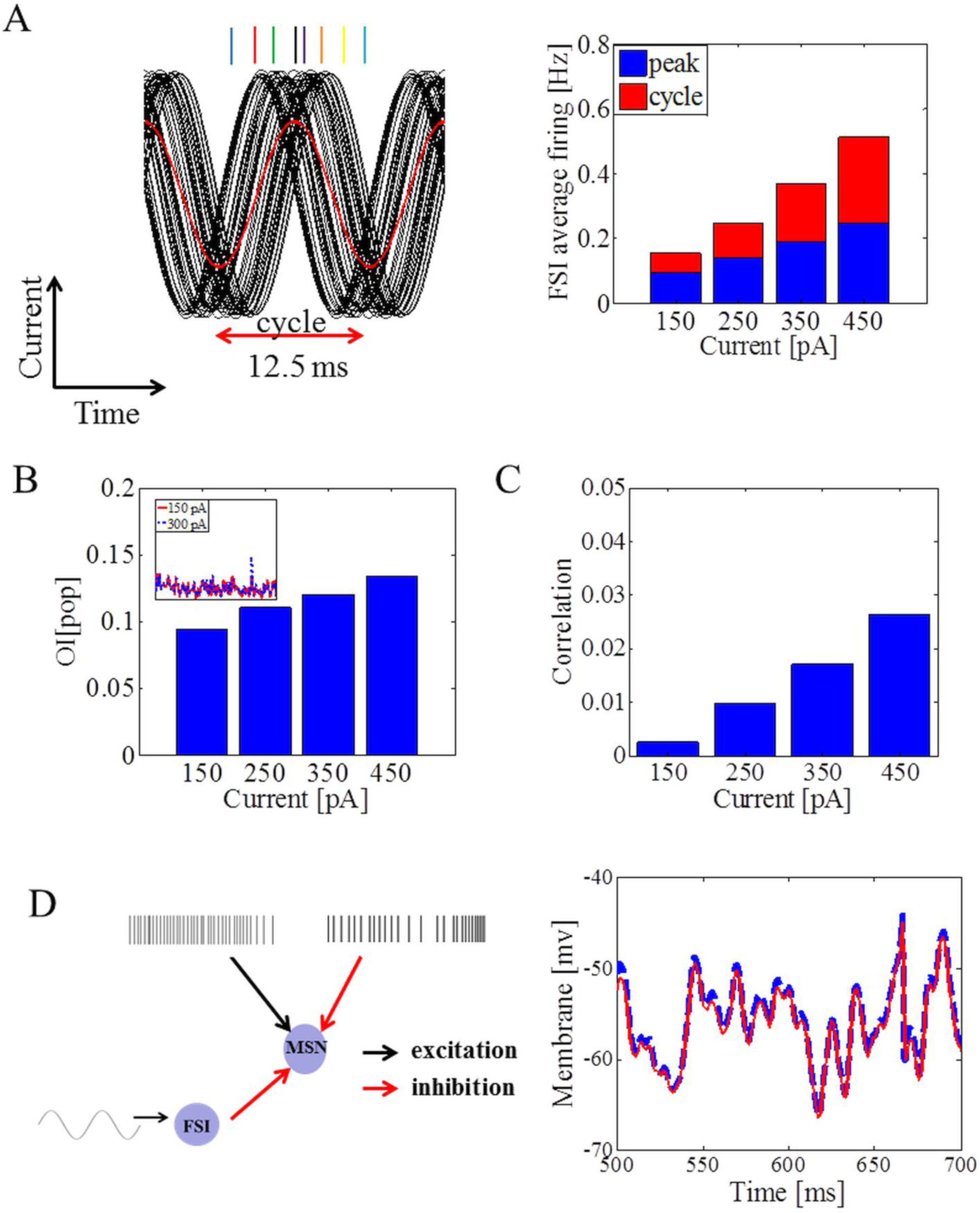
Firing of FSIs within an oscillation cycle and its influence on the emergence of oscillations in the MSN population. (A) The relationship between oscillatory inputs onto FSIs (black lines) and the timing of their overall spiking activity (left panel). Red line corresponds to the averaged value over all oscillatory inputs and the duration of one cycle of oscillations was 12.5 ms. The average firing of FSIs was low per oscillation cycle (red bars; right panel) but increased with a rise in the maximal amplitude of the oscillatory inputs. The proportion of neurons (blue bars) that fired in relation to the individual peaks of each FSI separately (defined in the range over ±2 ms of the peak value during each cycle) also increased with an increase in the maximal input oscillatory amplitude, but their ratio to the number of neurons that were entrained to the particular cycle declined. (B) Oscillatory index of the MSN population defined as an integral over power in the range of ±5 Hz around the driving frequency showed a steady increase correlated with the higher values of the maximal oscillatory input current (for A_max_=150 pA peak was not visible, while for A_max_=300 pA it was prominent). (C) The average pairwise correlation of spiking activity for FSIs. The values increased with an increase in the maximal amplitude of the oscillatory input, and the peak was visible for values of correlation < 0.01. (D) The neuron was first excited with an excitatory Poisson input (600 Hz) and one more Poisson-type synaptic input (500 Hz) was added, mimicking inhibitory inputs from other MSNs. The peak amplitude of the conductance transient was set to a biologically realistic value of −0.5 nS. Membrane potential (right plot, blue curve) was constantly reduced after adding FF inhibition (11 oscillatory driven FSIs; right plot, red curve).

### 3.3 The parameter dependence of emergent oscillations in the MSN population

Next, we varied the strength of the FF or FB inhibition, while the strength of other connection type was fixed (Fig. 6A). The strength of coupling between FSIs and MSNs was one of the key parameters of the striatal network that determined the successful propagation of oscillatory inputs onto FSIs to the MSN population. Upon decreasing the strength of the FF inhibition (from −3nS to - 1nS) three times, the peak in the power spectrum of the spiking activity of the MSN population at driving frequency was completely abolished (Fig. 6A, left panel). The peak could be re-established by setting a connection probability from FSIs to MSNs to high, biologically unrealistic values (>0.5). By contrast, the strength of FB inhibition did not play any prominent role in the transfer of oscillations in the MSN population. When we completely removed FB inhibition, the stability of the oscillatory peak in the power spectrum of the MSN population was not influenced (Fig. 6A, right panel). The peak increased only slightly from 0.0512 to 0.0514 (p>0.5). We also wanted to identify a minimum set of nodes that were sufficient to receive oscillatory input and transfer it to the MSN population. Our results showed that not all interneurons had to receive oscillatory inputs in order to successfully propagate it to MSN population (Fig. 6B, left panel). The maximum amplitude of the oscillatory input (A_max_) was equal to 350 pA (mean ± std firing of FSIs was equal to 30.98 ± 4.2 Hz) and was kept constant throughout all simulations, and the minimum set of FSIs sufficient to transfer oscillations onto MSNs depended on A_max_. In this scenario, when FSIs fired moderately, around half of the FSI population was sufficient to successfully transmit the received oscillatory inputs. Finally, the ongoing activity in the rest of the FSI network (different from the subset that received oscillatory input) could influence the survival of the peak in the MSN population. Fig. 6B (right panel) shows that when 36 FSIs received oscillatory input and Poisson background activity (see Materials and Methods), and the rest of the FSIs received Poisson drive that made them fire at two different frequencies, ~20 Hz and ~80 Hz, respectively, in the latter case the peak had a reduced value (0.033 vs 0.028, p=0.017).

**Figure 6:**
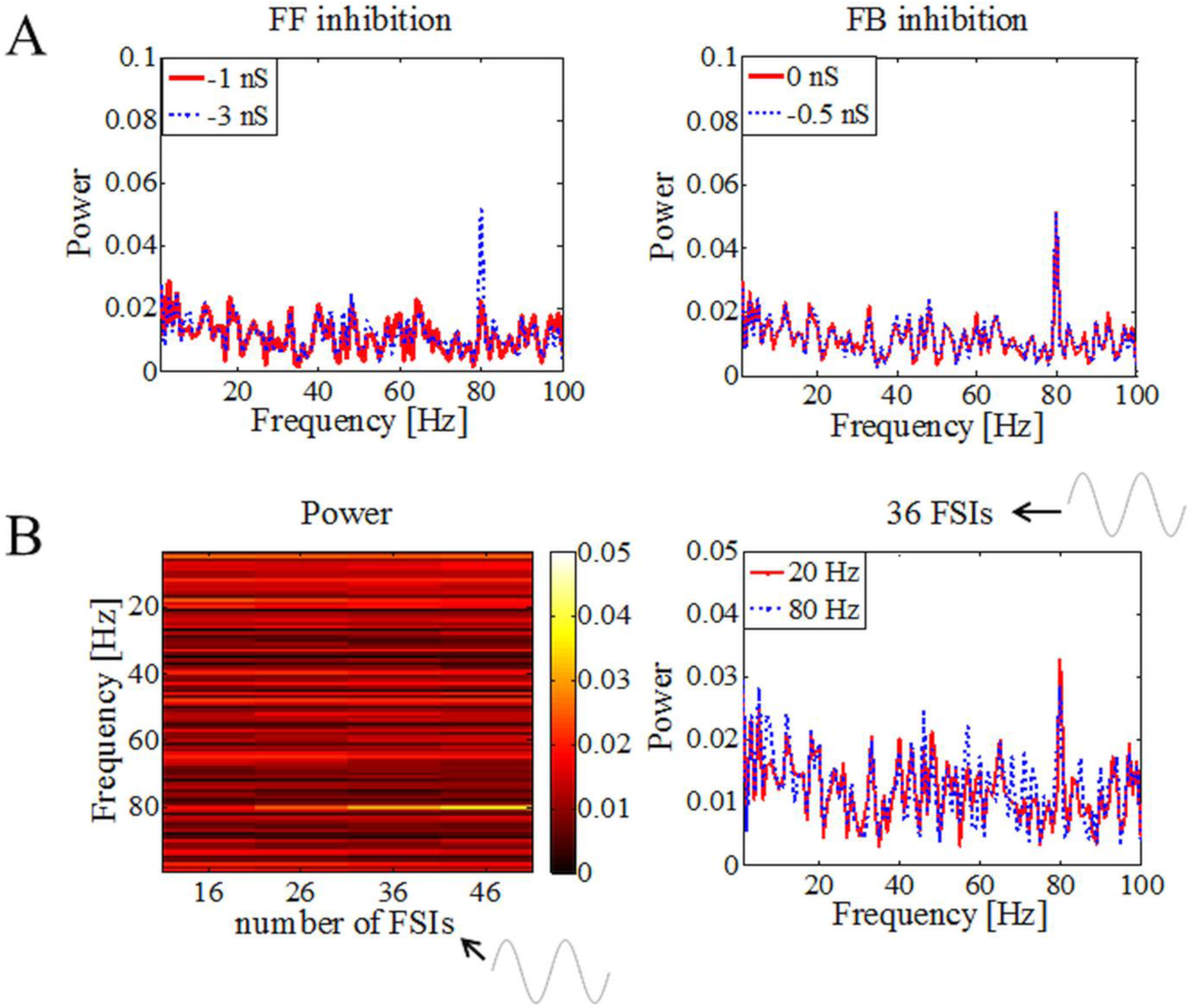
Effect of the network state on the emergence of oscillations in the MSN population. (A) Dependence of the strengths of the FF inhibition (left panel) and FB inhibition (right panel) on emergence of the oscillation frequency in the MSN population. (B) Dependence of the number of FSIs that are driven by asynchronous oscillatory inputs (left panel) and ongoing activity in the rest of FSI network (right panel) on the emergence of oscillations in the MSN population. Maximal amplitude of the oscillatory input was equal to 350 pA (mean ± std firing of FSIs was equal to 30.98 ± 4.2 Hz) and was kept constant throughout all simulations.

### 3.4 Interplay between the sole MSN network and asynchronous oscillatory drive

Finally, we wanted to see how selective stimulation of only MSNs with oscillatory currents (A_max_= 250 pA) lead to the emergence of oscillations. First, the stimulation was partial and only a portion of MSNs received oscillatory and Poisson-type synaptic background inputs while the rest of the MSN network received only Poisson-type synaptic background inputs and fired less than 1 Hz on average (see Materials and Methods). For 6.25 *%* of the oscillatory driven MSN population, the peak at driving frequency was not visible, while for 12.5% it was strong and stable. Next, we studied how the propagation of oscillatory activity depended on ongoing activity in the rest of the network (the MSNs that received only Poisson inputs). Our results showed that the entrainment of the MSNs to external oscillations was strongly influenced by the ongoing activity in the rest of the network in the way that it can destabilize the dynamics of the oscillatory propagation. When we increased the average firing of the rest of MSN network from <1 Hz to ~6 Hz, the oscillations at the input driving frequency completely disappeared (Fig. 7A). The spiking activity of the oscillatory driven population declined (average firing rate decreased from 1.65 Hz to 1.29 Hz) and it was completely masked by the high firing of the rest of the MSNs (those that were not oscillatory driven). The oscillations could reappear by increasing the firing of the oscillatory driven population to be close to the average firing of the rest of the MSN population. Crucially, in neither of these scenarios, the oscillatory driven population was able to transfer oscillations to the non-oscillatory driven population. Further we split MSNs into two populations. All neurons received Poisson background inputs and, additionally, the first half of MSNs received oscillatory inputs with the driving frequency set to 20 Hz, and the other half also received oscillatory inputs but with the driving frequency set to 30 Hz (in order to avoid interference of the first and second harmonics). We referred to the first half of the MSN population as the source network and the second half as the target network. Here, we investigated how the source network affected the oscillatory activity in the target network. Fig. 7B shows that the activity of the source network can impose the oscillatory activity in the target network, depending on the firing profile of the source network. When the average firing in the source was ~ 3.6 Hz, the source network could not impose oscillatory activity at its driving frequency onto the target network (Fig. 7B, blue line). We found that when the source network was in a state of high activity, a statistically significant peak was present in the power spectrum of the target network at the oscillatory frequency of the source network (Fig. 7B, red line; 0.0192 vs 0.0752, p<0.01). Similar results were obtained when the target network was only driven with Poisson inputs. Oscillatory input into the target network was kept constant in both cases (A_max_=160 pA) and we did observe a small peak decrease at the driving frequency of the target network in the second scenario (0.2218 vs 0.2134).

**Figure 7:**
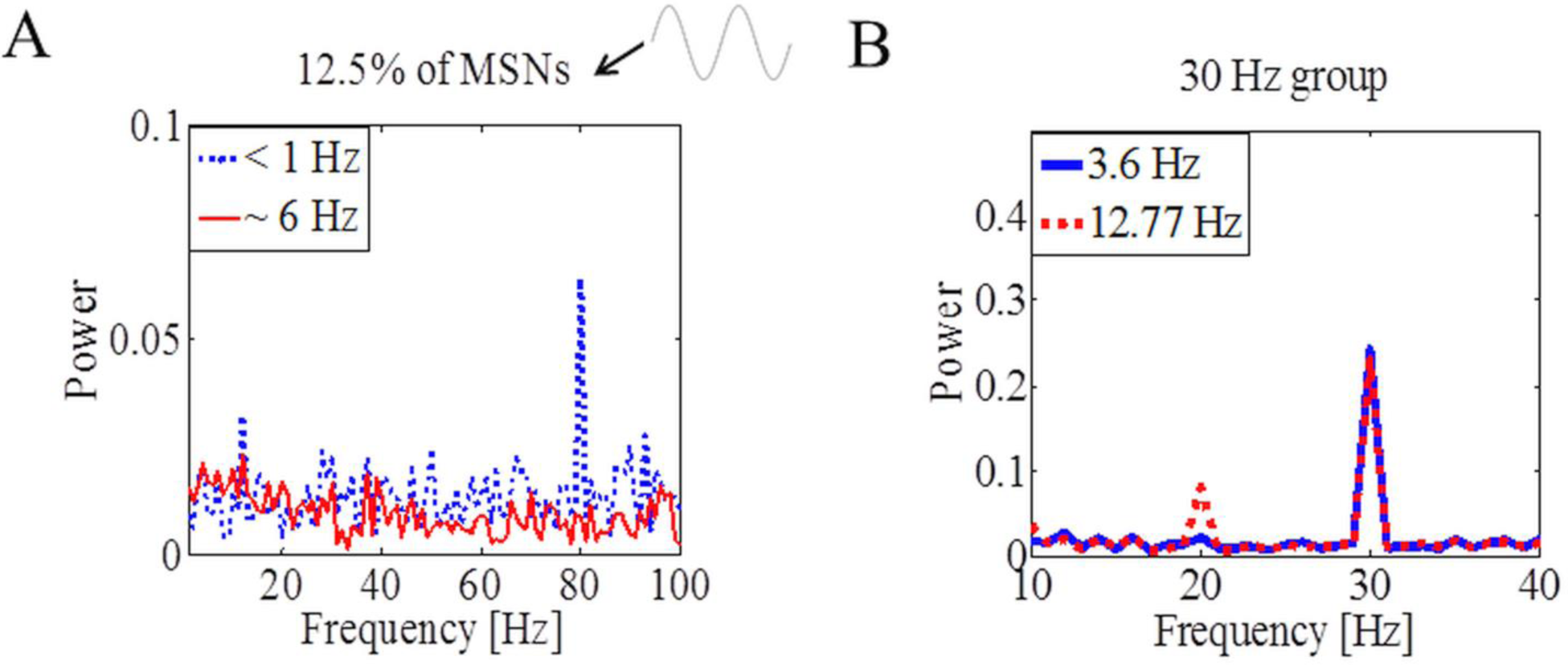
Response of sole MSN network to asynchronous oscillatory inputs. (A) The ongoing activity in the rest of the network can destabilize the dynamics of the oscillatory propagation. Increased average firing in the rest of the MSN network from <1 Hz to ~6 Hz, abolished the oscillations at the input driving frequency in the MSN network. (B) The MSN population was split into two equal sized populations. All neurons received Poisson background inputs and, additionally, the first population (referred to as the source network) received asynchronous oscillatory inputs with the driving frequency set at 20 Hz, and the other population (referred to as the target network) also received asynchronous oscillatory inputs but with the driving frequency set at 30 Hz. For the high activity state of the source network, a statistically significant peak was present in the power spectrum of the target network at the oscillatory frequency of the source network (0.0192 vs 0.0752, p<0.01).

## Discussion

Computational models of the striatum have started to reveal how cortical firing rates and correlations affect the response of striatal neurons [44–45]. The striatum is the largest component of the basal ganglia and its inhibitory circuits are known to be crucial for normal motor function [4, 5, 20, 21, 40]. A strong source of GABAergic inhibition to MSNs is FF inhibition from FSIs, although they comprise roughly only 1-2% of all striatal neurons [26, 31–33]. In contrast, the recurrent inhibition by MSNs (which represent ~95% of all striatal neurons) consists of many weak inputs [27, 28, 33, 51]. Striatal FSIs are scattered throughout the structure and they are also characterized by high but uncorrelated firing [36]. FSIs exist in other brain regions as well, such as the neocortex and hippocampus, and the experimental analysis showed that synapses among these neurons are highly specialized [58]. Several lines of evidence indicate the importance of the synchronous activity of FSIs for the generation of high frequency cortical rhythms, by generating a narrow window for effective excitation [58–60].

Oscillatory activity plays an important role in neural processing in many brain regions, including the basal ganglia [13, 21, 38–40, 57]. The striatal neurons receive massive direct inputs from the cortex, and FSIs appear to be more sensitive to the cortical inputs than MSNs [30, 61, 62]. There are clear evidences in favor of coherent reactivation between the hippocampus and striatum and the hippocampus projects to striatal parts that also receive convergent cortical inputs what might be of crucial importance for optimal sensory motor integration of striatal inputs [63, 64]. Experimentally, by simultaneous recording of striatal and cortical LFPs, it has been shown that oscillatory cortical inputs influence striatal activity in both rodents and primates, respectively [21, 40, 57, 65–67]. There are also peaks in gamma frequency range and recent experimental work has shown that striatal neurons respond to input-frequency components in cell-type-specific ways. Of all striatal cell types, only FSIs fired in relation to gamma frequencies where the MSNs’ frequency preference was not fixed [43]. Still, little is known about how high frequency oscillatory inputs propagate through striatal circuits and what role specific striatal cell types play in it. We developed a mathematical network model that encompassed neurons under the dendritic arborisation of a MSN to elucidate mapping between oscillatory stimulation parameters and ongoing network states on the MSN population’s oscillations. We have shown that the selective activation of a small portion of striatal FSIs via asynchronous oscillatory inputs could successfully entrain the MSN population to oscillate at the external input driving frequency. Although the firing of FSIs was uncorrelated, what really mattered was the sequence of activity of just a few FSIs relative to the specific oscillation cycle, allowing temporal summation of the inhibition exerted by these cells onto the MSNs. Besides imposing oscillatory rhythms onto MSNs, FSIs also induced fluctuations in oscillation power over time. FSIs often fire at high rates, but we showed that they might have a critical role in the propagation of high frequency oscillations in the MSN population even if their activity is kept low due to the strong inhibitory effects that FSIs can impose onto MSNs and the significant amount of shared connectivity. Further, we demonstrated that the features of the induced oscillations are influenced by internal states, especially ongoing activity. Therefore, our results support the idea of the precise orchestration of FSI activity that plays a key role in determining the pattern of the firing of MSNs, what might provide optimal integration of external inputs into striatal network. Also, one oscillatory MSN population could impose its rhythms onto other MSN population if it fired high, although MSNs generally fire very low, and if a large number of striatal neurons were oscillatory driven in the source network. Therefore, we suggest that the propagation of oscillatory inputs into the MSN population is efficient, if indirect, through FSIs. Recent study [68] investigated the role of striatal cholinergic interneurons in generating gamma oscillations in cortical-striatal circuits and reported that selective stimulation of cholinergic interneurons via optogenetics in normal mice amplified gamma oscillations. The study also demonstrated that gamma oscillations were largely restricted to the striatum what is out of scope of this paper where we were particularly interested in elucidating propagation of high frequency oscillations rather than local emergence. Also a previous study [43] showed that striatal cholinergic neurons fire in relation to the delta band frequencies. It is known that cholinergic neurons reduce the strength of the FSI synapses onto the MSNs by up to 13% and that cholinergic neurons’ agonists depolarize FSI [69]. Modulatory effects of cholinergic and dopaminergic neurons were taken into account by changing the effective strength of the FB and FF inhibitions what can shift certain thresholds (e.g. slight change of the firing threshold of FSIs when oscillations start to be present in the MSN population) but does not interfere with our main conclusions.

This work stands as the basis for future studies where one might use more detailed neuron models to study the influence of dendritic morphology and ion channel compositions on striatal oscillations. Overall, we have shown that a minor FSI population together with input-driven mechanisms might be responsible for the nature of spectral characteristics in the striatal network, thus enabling new perspectives on pharmacological intervention.

## Acknowledgements

The research leading to these results has received funding from the European Union Seventh Framework Programme (FP7/2007-2013) under grant agreement n°604102 (HBP), the Swedish Research Council, NIAAA (grant 2R01AA016022), Swedish e-Science Research Centre, and EuroSPIN – an Erasmus Mundus Joint Doctorate program. Any opinions, findings, and conclusions or recommendations expressed in this material are those of the authors and do not necessarily reflect the views of the funding organizations. The funders had no role in study design, data collection and analysis, decision to publish, or preparation of the manuscript.

## Author Contributions

Conceived and designed the experiments: J.B., A.K., and J.H.K. Performed the experiments: J.B. Analysed the data: J.B., A.K., and J.H.K. Contributed to the writing of the manuscript: J.B., A.K., and J.H.K.

## Data Availability

All relevant data are within the paper.

